# Unraveling Dengue Serotype 3 Transmission in Brazil: Evidence for Multiple Introductions of the 3III_B.3.2 Lineage

**DOI:** 10.1101/2025.01.23.634513

**Authors:** James Siqueira Pereira, Svetoslav Nanev Slavov, Isabela Carvalho Brcko, Gabriela Ribeiro, Vinicius Carius de Souza, Igor Santana Ribeiro, Iago Trezena T. De Lima, Gleissy Adriane Lima Borges, Katia Cristina de Lima Furtado, Shirley Moreira da Silva Chagas, Patricia Miriam Sayuri Sato Barros da Costa, Talita Adelino, Felipe C. de Melo Iani, Luiz Carlos Junior Alcantara, Verity Hill, Nathan D. Grubaugh, Sandra Coccuzzo Sampaio, Maria Carolina Elias, Marta Giovanetti, Alex Ranieri J. Lima

**Affiliations:** Centro para Vigilância Viral e Avaliação Sorológica (CeVIVAS), Instituto Butantan, Av. Vital Brazil, 1500, São Paulo, SP, 05503-900, Brazil; Programa Interunidades de Pós-graduação em Bioinformática, Universidade de São Paulo, Av. Prof. Lineu Prestes, 748, São Paulo, SP, 05508-900, Brazil; Laboratório Central de Saúde Pública do Estado do Pará (LACEN-PA), Av. Almirante Barroso, 4341, Belém, PA, 66613-710, Brazil; Laboratório Central de Saúde Pública do Estado de Minas Gerais (LACEN-MG), Fundação Ezequiel Dias, Rua Conde Pereira Carneiro, 80, Belo Horizonte, MG, 30510-010, Brazil; Instituto Rene Rachou, Fundação Oswaldo Cruz, Av. Augusto de Lima, 1715 - Barro Preto, Belo Horizonte - MG, 30190-002, Brazil; Department of Epidemiology of Microbial Diseases, Yale School of Public Health, New Haven, CT 06510, USA; Department of Sciences and Technologies for Sustainable Development and One Health, Università Campus Bio-Medico di Roma, Via Álvaro del Portillo, 21, Rome, RM, 00128, Italy; Oswaldo Cruz Institute, Oswaldo Cruz Foundation, Av. Brasil, 4365, Manguinhos, Rio de Janeiro, RJ, 21040-900, Brazil

**Keywords:** Dengue lineage, phylogeography, outbreak, genomic surveillance

## Abstract

Dengue, caused by DENV 1-4, remains a global public health concern, with Brazil experiencing some of the largest epidemics. The reemergence of DENV-3 in Brazil between 2023 and 2024 has raised concerns about new outbreaks due to the absence of sustained circulation of this serotype in recent years. This study investigates the dynamics of DENV-3 in Brazil, focusing on the spread of the 3III_B.3.2 lineage within genotype 3III and its introduction routes. We analyzed 1,536 DENV-3 genomes, all classified as genotype 3III, the dominant DENV-3 genotype in Brazil since 2001. Phylogenetic analysis identified the 3III_B.3.2 lineage in all recent Brazilian cases, with detections also reported in Central America, the United States, and Europe. At least six independent introduction events of this lineage into Brazil were identified, with the Caribbean region and Costa Rica as the primary sources. The earliest introduction likely occurred in late 2022 in Roraima, followed by introductions in São Paulo, Minas Gerais, and Pará. While one instance of interstate transmission was detected — from São Paulo to Minas Gerais — our findings indicate that external introductions, rather than domestic spread, were the primary drivers of DENV-3 circulation during this period. These results underscore the importance of continued genomic surveillance and coordinated public health strategies to monitor and mitigate future outbreaks.

## 1. Introduction

Dengue fever, the most prevalent arbovirus disease worldwide, is caused by dengue virus (DENV), a member of the *Flaviviridae* family (genus *Orthoflavivirus, Orthoflavivirus dengue*, DENV)[1]. The main transmission route for DENV is through the bites of female mosquitoes from the *Aedes* genus, particularly *Aedes aegypti* and *Aedes albopictus*. However, other transmission routes, such as blood transfusion[2,3] and vertical transmission[4] have also been observed. Clinically, the infection ranges from mild to severe, with the latter manifesting as hypovolemic shock due to plasma leakage, which is also associated with high mortality. The reason why some patients develop severe forms of DENV is not fully understood, but the presence of heterologic antibodies to other subtypes has been suggested as a primary factor[5].

DENV is classified into four distinct serotypes (DENV-1 to DENV-4), which are phylogenetically related[6]. Moreover, within each serotype, there is genetic variability, with multiple genotypes identified. Brazil, a global leader in the number of DENV cases, reported over 9.1 million cases in 2024 alone, with significantly increased mortality from hemorrhagic dengue fever (5,008 deaths)[7]. The country is also hyperendemic for DENV, with the circulation of the four serotypes, although recent outbreaks have been mainly caused by serotypes 1 and 2 with sporadic records of the other two serotypes[8]

DENV-3 is classified into five distinct genotypes (I to V), with genotype III being the most widespread and associated with major outbreaks in Asia, Africa, and the Americas[9]. In Brazil, the first autochthonous transmission of this genotype was reported in the late 2000s, marking the introduction of this serotype in the country[10]. Recent reports have indicated a notable increase in the detection of DENV-3, suggesting a resurgence of this serotype in frequency and geographic distribution[11–13]. This situation poses a significant risk for the emergence of new outbreaks, particularly given the historical absence of sustained DENV-3 circulation and the probable low serological coverage.

By epidemiological week 26 in 2024, 409 cases of DENV-3 were confirmed in Brazil, with the highest detection rates in the Northern and Southeastern regions[8]. In this context, our study investigates the recent spread of DENV-3 in Brazil using a phylogeographic approach, with a focus on the classification of circulating lineages and the detection of eleven new cases in the Northern and Southeastern regions of the country.

## 2. Materials and Methods

### 2.1 Clinical Sample Collection and Genome Sequencing

#### 2.1.1 The CeVIVAS initiative

The samples used in this study were provided by the CeVIVAS (Center for Viral Surveillance and Serological Assessment), a collaborative initiative spearheaded by Instituto Butantan in partnership with public health laboratories and institutions. CeVIVAS aims to strengthen viral genomic surveillance and serological assessment efforts in the country (https://bv.fapesp.br/en/auxilios/110575/continuous-improvement-of-vaccines-center-for-viral-surveillance-and-serological-assessment-cevivas/enter-for-viral-surveillance-and-serological-assessment-cevivas/). This study was approved by the Institutional Ethics Committee of the Faculty of Medicine of the University of São Paulo (Decision number CAAE: 68586623.0.0000.0068)

#### 2.1.2 Clinical samples

The samples analyzed in this study were collected between epidemiological weeks 4 and 44 on LACEN Pará (n=2) and LACEN Minas Gerais (n=9), and were confirmed as DENV-3, using the CDC-established protocol[14], exhibiting Ct values below 30. The RNA extraction was performed to all samples using the Extracta kit AN viral (Loccus) in an automated extractor (EXTRACTA 96, Loccus) according to the manufacturers’ specifications. The confirmation of viral RNA was performed using multiplex TaqMan real-time PCR detecting all four serotypes and targeting the 3’UTR genomic portion. DENV sequencing was performed using primer sets (Table A.1) specific for DENV-3. Genomic libraries were prepared using the COVIDSeq protocol, following the manufacturer’s instructions, except for the used primers. The pooled libraries were normalized to 4 nM, denatured with 0·2 N NaOH and 400 mM Tris-HCl (pH 8), and sequenced using the NextSeq 500/550 High Output Kit v2.5 (300 cycles) on the Illumina NextSeq 2000.

### 2.2 Genome Assembly Pipeline

The assembly pipeline employed, detailed in Fig. A.1, is part of the VIPER (Viral Identification Pipeline for Emergency Response), which is publicly accessible at https://github.com/V-GEN-Lab/viper. Comprehensive descriptions of the specific parameters and software utilized are provided in Appendix 1 (p. 4).

### 2.3 Classification of DENV-3 genomes and phylogenomic reconstruction

We selected all complete DENV-3 genomes (genome size >10 Kb) collected from human hosts and available in GenBank, using the NCBI Virus portal (https://www.ncbi.nlm.nih.gov/labs/virus/vssi/#/), accessed on August 22, 2024. Additionally, we retrieved all Brazilian complete genomes (>10 kb) available in GISAID EpiArbo[15], collected from January 1, 2022, onward, with complete collection date information. The dataset (EPI_SET_250123aq) (10.55876/gis8.250123aq) was restricted to sequences associated with publications or deposited by laboratories collaborating with the CeVIVAS project (see Disclaimer). The final dataset (n=2,358), along with the eleven genomes generated in this study, was classified using Nextclade[16], with the DENV-3 reference dataset, following the new Dengue Lineages nomenclature[17]. All genomes belonging to genotype III were selected, as all Brazilian sequences belonged to this genotype.

The dataset of DENV-3 genotype 3III (n=1,536; metadata available at Zenodo https://doi.org/10.5281/zenodo.13885992 in Supplementary File 1) was aligned using the Augur align tool from the Augur program toolkit v24.1.0[18], with the MAFFT v7.520 algorithm[19]. A maximum likelihood (ML) phylogenetic tree was then constructed using IQ-TREE v2.2.6, applying the GTR+F+I+G4 nucleotide substitution model, as determined by ModelFinder within IQ-TREE2. The robustness of the tree topology was evaluated through 10,000 UFBoot replicates.

### 2.4 Spatiotemporal and viral recent spread reconstruction of DENV-3

To investigate the recent spread of DENV-3 in Brazil, we employed a discrete Bayesian phylogeographic inference approach using BEAST v1.10.4[20]. To optimize computational efficiency, branches containing recent Brazilian samples (collected from 2022 onwards) were identified from the Maximum Likelihood (ML) phylogenetic tree, and the corresponding sequences were retrieved (n=88). Outlier sequences (n=4), with temporal values deviating by more than 1.5 interquartile ranges in root-to-tip regression analysis, identified using TempEst v1.5.3[21], were excluded[12]. Time-scaled phylogenetic trees were inferred using a relaxed molecular clock model[22]. Ancestral states were reconstructed based on a continuous-time Markov chain (CTMC) prior[23] with discrete spatial diffusion, using an asymmetric substitution model to infer migration events[24]. Bayesian stochastic search variable selection (BSSVS) was implemented to identify statistically supported migration routes. Two independent Markov Chain Monte Carlo (MCMC) chains were run for 200 million generations each. Convergence was evaluated by calculating the Effective Sample Size (ESS >200) for all parameters using Tracer v1.7.1[25]. The Log files are available at Zenodo https://doi.org/10.5281/zenodo.13885992 in Supplementary Files 2 and 3. After discarding the initial 10% as burn-in, the maximum clade credibility (MCC) tree was summarized with TreeAnnotator[26]. The final time-scaled phylogeny was visualized in FigTree v1.4.5[27], and spatiotemporal information embedded in the posterior distribution of trees was extracted and mapped using the R package ‘seraphim’ (version 1.0)[28]. The metadata for the final dataset is available in Table A.2.

## 3. Results

The 11 genomes generated in this study were obtained from samples collected between epidemiological weeks 4 and 44 of 2024. The samples, confirmed as DENV-3 (see Methods for details), exhibited Ct values below 30, with genomic coverages ranging from 65,7% to 91,7%, resulting in an overall mean coverage of 79.62%.

Of the 2,376 DENV-3 sequences analyzed, 1,537 were classified as genotype 3III, the only genotype detected in Brazil, except for a single sample collected in 2006 (GenBank accession OQ727062). Our analysis reclassified this sample as genotype V, as it was originally labeled as genotype I in the GenBank database. Among the three major lineages within genotype 3III, only 3III_B and 3III_C were identified in Brazil, each circulating during distinct periods (Figure 1A). Lineage 3III_C was detected exclusively until 2009, whereas lineage 3III_B, specifically sub-lineage 3III_B.3.2, has been circulating since May 2022 in countries such as Angola, Cuba, the Dominican Republic, Italy, Nigeria, Suriname, and the United States (Fig. A.2). This sub-lineage has been associated with all Brazilian genomes collected since 2023 (n = 39), including the eleven genomes sequenced in this study.

**Figure 1.**
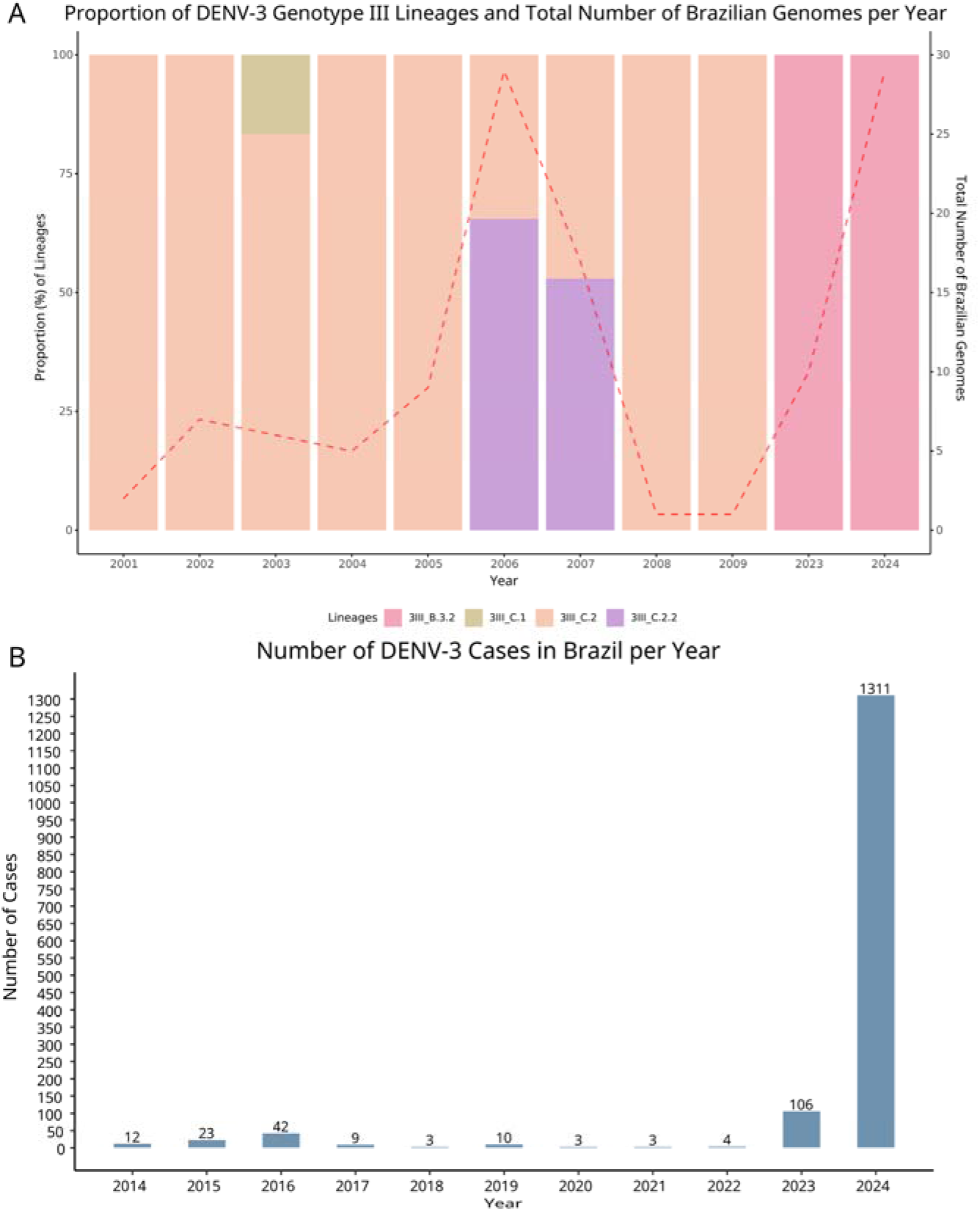
(A) Proportion of DENV-3 genotype III lineages detected in Brazil from 2001 to 2024. The left Y-axis shows the percentage of each lineage (colored bars) relative to the total number of sequenced samples per year (X-axis). The colors represent specific lineages: 3III_B.3.2 (pink), 3III_C.1 (green), 3III_C.2 (peach), and 3III_C.2.2 (lavender). The right Y-axis displays the total number of Brazilian genomes sequenced each year, represented by the red dashed line. The absence of bars for certain years (2010–2022) reflects a lack of available genomic data for these periods. (B) Annual number of DENV-3 cases in Brazil from 2014 to 2024. The Y-axis on the left indicates the number of reported cases, and the X-axis shows the years (data sourced from the Brazilian Notifiable Diseases Information System - SINAN [TabNet, http://tabnet.datasus.gov.br/cgi/deftohtm.exe?sinannet/cnv/denguebbr.def], accessed on 08 January 2025, and available on Zenodo at https://doi.org/10.5281/zenodo.13885992 in Supplementary File 4).

In 2023, the number of reported DENV-3 infections in Brazil showed a significant increase, rising from four cases in 2022 to 106 cases. This upward trend continued into 2024, with 1,311 cases reported (Figure 1B). São Paulo recorded the highest number of cases, reporting 510 probable infections, followed by Minas Gerais with 302 cases. Among the 22 Brazilian states listed in the case notification system, 17 reported at least one DENV-3 case in 2024.

Phylodynamic analyses of genotype 3III sequences, particularly within lineage 3III_B.3.2, revealed a close evolutionary relationship between Brazilian sequences and those from the Caribbean region, including two sequences from Italy (Fig. A.3), as previously suggested[29,30]. To investigate the dispersion patterns of the 3III_B.3.2 lineage in Brazil and its potential introduction routes, we performed a phylogeographic analysis incorporating all Brazilian samples alongside the most closely related sequences from Cuba, USA, Costa Rica, Dominican Republic, Italy, Puerto Rico, Suriname, and Haiti. This dataset (n=84) had a strong temporal signal (Figure 2A) and provided evidence of multiple independent introductions of the 3III_B.3.2 lineage into Brazil. The earliest introduction event probably occurred in late October 2022 (Figure 2C and Figure 3), likely originating from Cuba and entering the state of Roraima, in Northern Brazil. Additionally, six other introduction events were identified across several Brazilian states from the Caribbean region and Costa Rica. The most recent introduction of the 3III_B.3.2 lineage into the state of Pará was estimated to have occurred in late February 2024, originating from Cuba.

**Figure 2.**
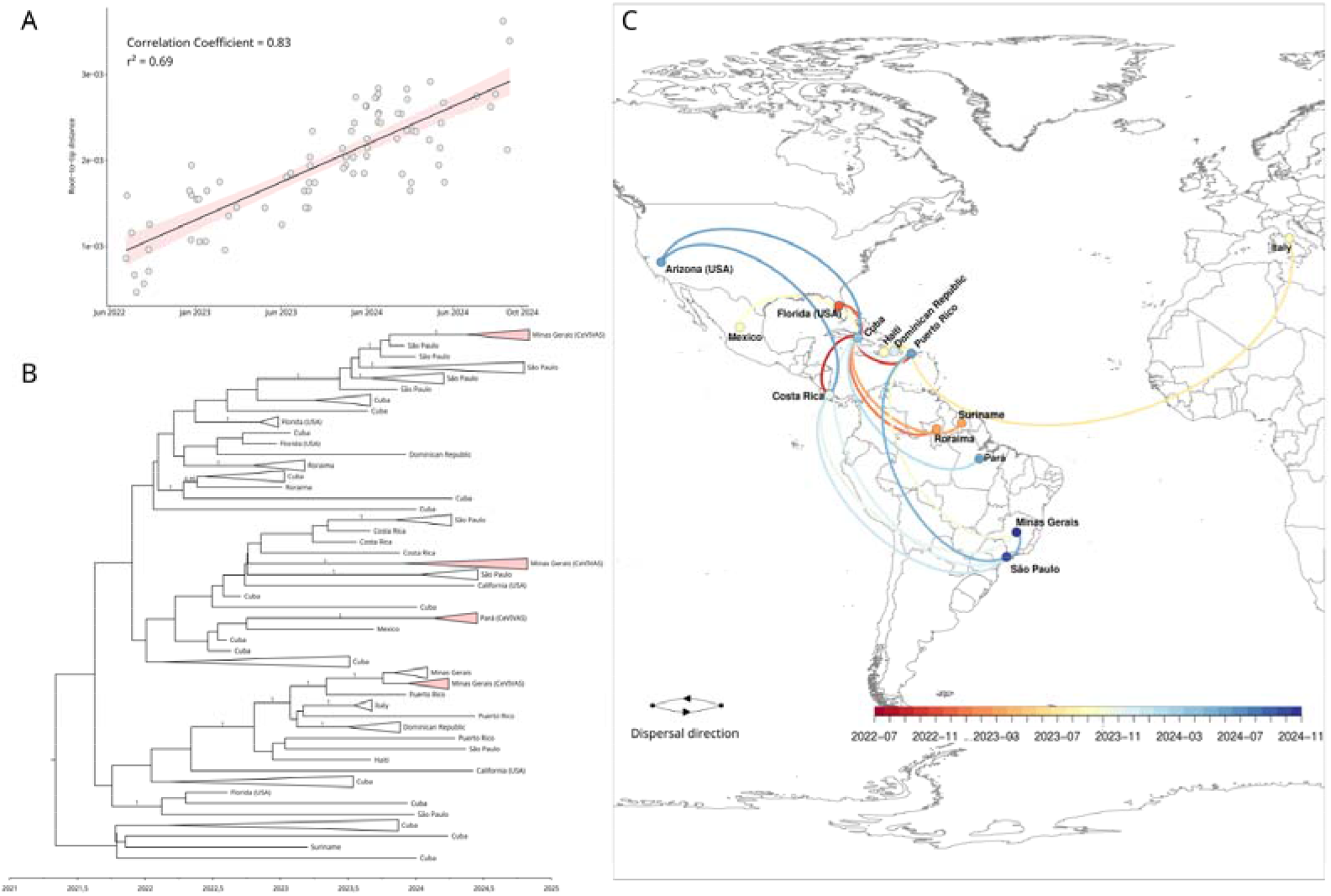
Phylogenetic analysis and geographical distribution of the 3III_B.3.2 lineage from 2021 to 2024. (A) Linear regression between genetic distance and time (in years) shows a significant correlation (correlation coefficient = 0.83, r^2^ = 0.69), indicating the temporal evolution of the virus. (B) Maximum Clade Credibility (MCC) tree based on genomic sequences from Brazil, with the addition of sequences from Cuba, USA, Dominican Republic, Costa Rica, Mexico, Puerto Rico, Italy, Haiti, and Suriname. The posterior probability values for branches associated with introduction and dispersal events in Brazil are also indicated on the tree. The shaded (in red) triangle highlights the genomes generated in this study. (C) Geographic map showing the spread of the analyzed sequences. The arrows indicate the dispersal direction of the virus over time, from 2022 to 2024. The color of the lines, and the circles, corresponds to the time, as shown in the legend, ranging from the earliest in blue to the most recent in green and the direction of the arrows represent the dispersal direction. A dynamic geographic map, generated using spread.gl[31], is available as an HTML file Zenodo https://doi.org/10.5281/zenodo.13885992 in Supplementary File 5.

**Figure 3.**
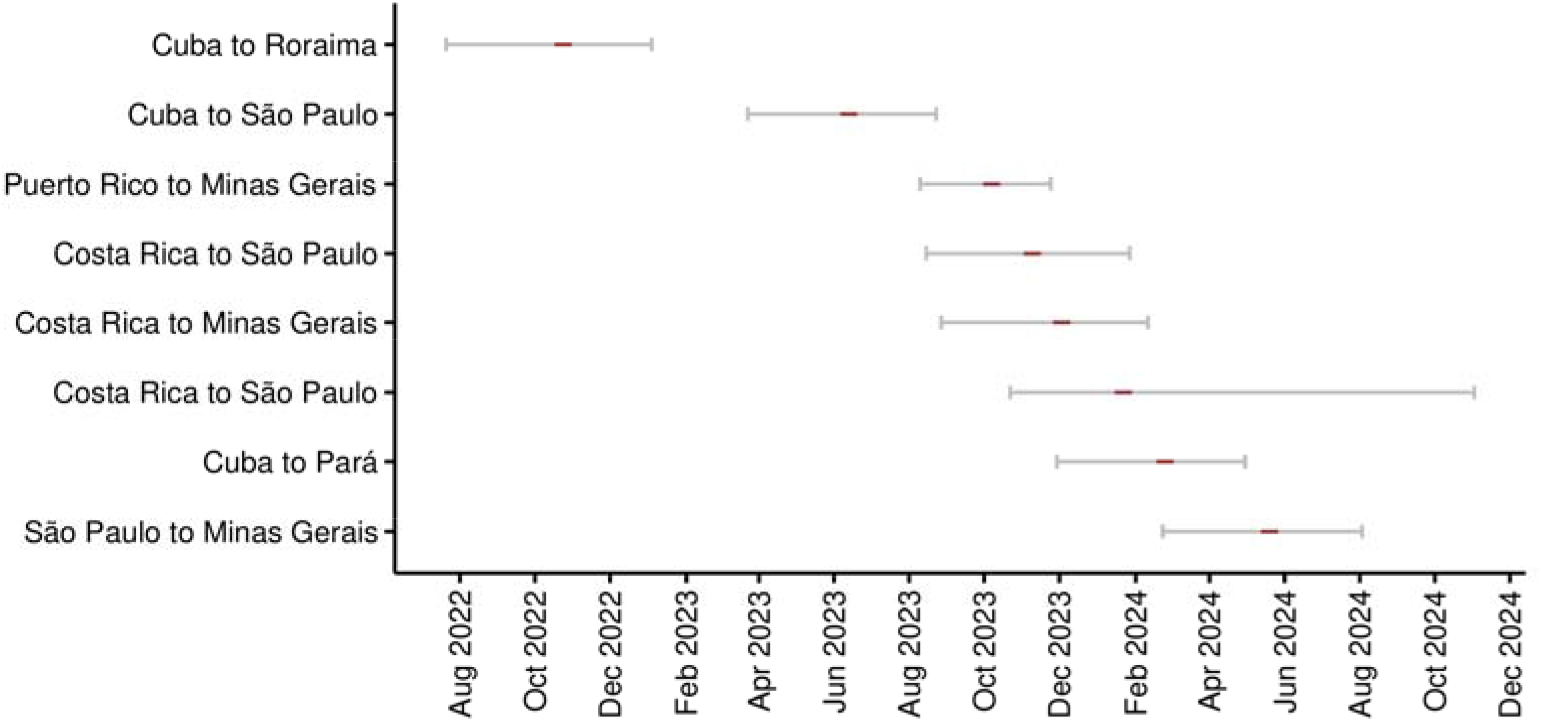
Temporal dynamics of the introductions and spread of lineage 3III_B.3.2 in Brazil. Red represents temporal estimates for the most recent common ancestor (TMRCA) of each lineage introduction, while the associated uncertainty (95% HPD interval) is shown by horizontal error bars.

Two sequences from the state of São Paulo, Southeast Brazil, with collection dates of 25 December 2023 and 5 May 2024, were located on distinct branches of the MCC tree, each tracing back to different ancestral nodes from Cuba and Puerto Rico. However, this inference remains uncertain due to the low posterior probability of these nodes in the phylogenetic tree. We also identified one probable internal dispersal event of the 3III_B.3.2 lineage within Brazil: from São Paulo to Minas Gerais (Southeast) around mid-May 2023.

## 4. Discussion

The 2023/2024 dengue epidemic in Brazil stands as one of the most severe in recent decades, predominantly driven by serotypes 1 and 2. However, the reemergence of DENV-3, after more than 13 years of limited circulation, introduced a new dynamic to the epidemic. The prolonged absence of serotype 3 likely left a substantial portion of the population seronegative for DENV-3, heightening the risk of large-scale outbreaks similar to those witnessed in 2002[32,33]. From 2014 to 2022, only 109 DENV-3 cases were reported in Brazil, with the peak occurring in 2016 (42 cases), as documented in the Brazilian Notifiable Diseases Information System (SINAN) database (see Supplementary File 4). This trend shifted dramatically in 2023, with cases surging to 106, signifying the virus’s resurgence, with detections in Roraima (Northern), Paraná (Southern), and Minas Gerais (Southeast)[12,13].

Historically, DENV-3 sequences from Brazil up until 2009 were primarily classified as belonging to the 3III_C lineage, particularly sublineages 3III_C.2 and 3III_C.2.2. However, genomic surveillance from 2023 to 2024 revealed that the circulating viral population had shifted entirely to lineage 3III_B.3.2. This replacement of viral lineages is significant, as such shifts are often associated with altered outbreak dynamics and epidemiological patterns[34–36]. Despite the lack of genomic data from Brazil between 2010 and 2022, the detection of a limited number of DENV3 cases by serology from 2014 to 2022 suggests the residual circulation of lineage 3III_C. This hypothesis is supported by the increase in cases observed in 2023, which coincides with our phylogeographic data indicating the introduction of lineage 3III_B.3.2 lineage into Brazil around October 2022, originating from the Caribbean region. This finding aligns with a previous study suggesting that the lineage was introduced into the Caribbean at the end of 2020, with subsequent spread to Suriname, the United States, and eventually Brazil[11,12]. These introductions underscore the critical role of international dissemination in the resurgence of DENV-3, particularly in northern Brazil.

Our analyses indicate that, between 2022 and 2024, multiple independent introductions from the Caribbean region contributed to the resurgence of DENV-3 in various parts of Brazil. Although the number of cases had been relatively low in previous years, the virus re-emerged in 2023, with the highest incidence rates concentrated in the northern states[8]. Interestingly, we identified only a single instance of interstate transmission occurring in the Southeastern region of Brazil, between the states of São Paulo and Minas Gerais, likely occurring in mid-May 2024, suggesting that external introductions, rather than domestic spread, were the primary drivers of DENV-3 circulation during this period.

However, given the notable expansion of DENV transmission in Southern Brazil, particularly in the states of Santa Catarina and Rio Grande do Sul—driven by factors such as increased environmental suitability for Aedes, a susceptible population, and multiple independent viral introductions[37]—we infer that additional international introductions might have occurred. Furthermore, other domestic transmission routes may also play a significant role in the dissemination of this lineage.

Two of the new DENV-3 genomes generated in this study were collected on May 14 and June 9, 2024, from symptomatic individuals in a rural area of Altamira, Pará. These samples represent some of the first documented cases of DENV-3 in the state since 2018, as indicated by SINAN data (TabNet, http://tabnet.datasus.gov.br/cgi/deftohtm.exe?sinannet/cnv/denguebbr.def, accessed on January 8, 2025) and Supplementary File 6 (Zenodo: https://doi.org/10.5281/zenodo.13885992). Phylogeographical analysis suggests that the 3III_B.3.2 lineage was introduced into Pará from Cuba around early February 2024, highlighting international viral movement and prior undetected circulation in the region.

Additionally, nine of the new DENV-3 genomes were collected from Minas Gerais between epidemiological weeks 4 and 44 of 2024, reflecting sustained circulation of the 3III_B.3.2 lineage. That year, Minas Gerais recorded its highest number of DENV-3 cases in the past decade (n=302), according to SINAN data (Supplementary File 6). Phylogeographical analyses identified at least two international introduction events of this lineage into the state. The first occurred in early October 2023, likely originating from the Caribbean region, specifically Puerto Rico, as suggested by previous studies[12,13]. A second introduction from Costa Rica was detected in early December 2023. In addition to the domestic transmission observed, likely occurring in mid-May 2024.

Overall, our findings indicate that introductions of the 3III_B.3.2 lineage into different Brazilian states likely occurred two to three months before their detection, underscoring the challenges of detecting viral circulation in real-time. However, underreporting of cases and limited data availability (see Disclaimer) introduce significant biases that could place these introductions earlier than the data suggest.

## 5. Conclusion

Our study underscores the reemergence of DENV-3 in Brazil, driven predominantly by the introduction and circulation of the 3III_B.3.2 lineage. Multiple independent introductions from the Caribbean and Costa Rica, followed by subsequent internal spread within Brazil, highlight the dynamic patterns of viral migration. These findings emphasize the necessity for enhanced genomic surveillance and coordinated public health interventions to mitigate the potential for the emergence of widespread DENV-3 outbreaks, particularly in regions with limited prior exposure to this serotype. Ongoing surveillance efforts will therefore be critical in identifying emerging DENV lineages and guiding timely responses to prevent future DENV epidemics in Brazil.

## Acknowledgments

We are grateful to Dr. Marcio Roberto Teixeira Nunes for his valuable consultation during the analysis and insightful contributions throughout the study. We disclose that ChatGPT was employed only to review the clarity and use of the English language in this manuscript. We gratefully acknowledge all data contributors, i.e., the Authors and their Originating laboratories responsible for obtaining the specimens, and their Submitting laboratories for generating the genetic sequence and metadata and sharing via the GISAID Initiative, on which this research is based. This work used resources from the High-Performance Computing System of the Center for Bioinformatics and Computational Biology (NBBC) of the Butantan Institute

## Authors’ contributions

JSP and ARJL conceptualized the study and curated the data, as well as performed the formal analyses. SCS and MCE acquired the funding. JSP, GR, GALB, KCLF, PMSSBC, SMSC, TA, FCdMI, MG, and ARJL conducted the investigation. JSP, ICB, VCS, ISR, ITTL, VH, MG, and ARJL developed the methodology. ARJL managed the project, while GALB, KCLF, PMSSBC, SMSC, TA and FCdMI provided resources. JSP was responsible for data visualization. JSP, SNS, and ARJL wrote the original draft, and JSP, SNS, ICB, GR, VCS, VH, NDG, SCS, MCE, MG, and ARJL reviewed and edited the manuscript.

## Funding

This research was supported by FAPESP (São Paulo Research Foundation) under grant number 21/11944-6. JSP received PhD fellowship support through grants 88887.969077/2024-00 from CAPES (Fundação Coordenação de Aperfeiçoamento de Pessoal de Nível Superior) and 2023/11919-7 from FAPESP (Fundação de Amparo à Pesquisa do Estado de São Paulo.) MG is funded by PON “Ricerca e Innovazione” 2014-2020 and by the CRP-ICGEB RESEARCH GRANT 2020 Project CRP/BRA20-03, Contract CRP/20/03.

## Data Statement

The dataset of sequences retrieved from GISAID can be accessed using the EPI_SET ID: EPI_SET_250123aq (10.55876/gis8.250123aq). Additionally, all sequence accession numbers, associated metadata used in this study, BEAST analysis logs, publicly available serological data, and an .html file illustrating the introduction routes are available on Zenodo: https://doi.org/10.5281/zenodo.13885992.

## Conflict of Interest Statement

The authors declare no conflicts of interest related to this study.

## Disclaimer

The authors acknowledge the availability of additional DENV-3 sequences in the GISAID database. However, these could not be included in the study due to a lack of authorization from their submitters. While sampling bias is a recognized limitation, the inclusion of these sequences would primarily enhance insights into viral dissemination routes to other Brazilian states, without significantly impacting the study’s main findings.

## Notes

https://doi.org/10.5281/zenodo.13885992

### Competing Interest Statement

The authors have declared no competing interest.

