## Supplementary_material for "Unraveling Dengue Serotype 3 Transmission in Brazil: Evidence for Multiple Introductions of the 3III_B.3.2 Lineage"

### Extended Methods

**Supplementary Table 1. Set of DENV-3 virus-specific primers designed to generate genomes**

| **Primer Name** | **Primer pool no.** | **Sequence (5'-3')** |
|---|---|---|
| DENV3_1_LEFT | 1 | TCAATATGCTGAAACGCGTGAGA |
| DENV3_1_RIGHT | 1 | GCCTCGGTCTTCTGAAGCTCTA |
| DENV3_2_LEFT | 2 | GGGAGTAGGAAACAGAGATTTTGTGG |
| DENV3_2_RIGHT | 2 | GAGTATTGTCCCATGCTGCGTT |
| DENV3_3_LEFT | 1 | ACCAATAGAGGGAAAAGTGGTGC |
| DENV3_3_RIGHT | 1 | TGGCCTCGAACATCTTCCCAAT |
| DENV3_4_LEFT | 2 | GCTGAACCTCCTTTTGGGGAAA |
| DENV3_4_RIGHT | 2 | TTCCACCTCCCACACATTCCAT |
| DENV3_5_LEFT | 1 | GGCAAAAATAGTGACAGCTGAAACA |
| DENV3_5_RIGHT | 1 | TCTCCATGTTATTTGCCCTGAGAGA |
| DENV3_6_LEFT | 2 | AGGTGGACAACTTCACAATGGG |
| DENV3_6_RIGHT | 2 | CTCAGCCTCTTCCTCCCATGTT |
| DENV3_7_LEFT | 1 | GCCAGTCTTCGAGCATGAGGAA |
| DENV3_7_RIGHT | 1 | TTCCCAGGCTCTACGGCAATAA |
| DENV3_8_LEFT | 2 | AAAGACTGGAACCAAACTGGGC |
| DENV3_8_RIGHT | 2 | TGCCTGAATTCCATGAGCGTTC |
| DENV3_9_LEFT | 1 | GCAACAAAATCTGAACACACAGGA |
| DENV3_9_RIGHT | 1 | CCCTCCTCATGAGTTCCACGAA |
| DENV3_10_LEFT | 2 | AGACCATGCTCACTGGACAGAA |
| DENV3_10_RIGHT | 2 | TATGCGAGTTGGTTGTCTTGGG |
| DENV3_11_LEFT | 1 | GGAAAGACTTCAATAGGACTCATTTGTG |
| DENV3_11_RIGHT | 1 | CAGGTGATCCTTCCCAGAGTGT |
| DENV3_12_LEFT | 2 | GTGGATGGGATAATGACAATAGACCT |
| DENV3_12_RIGHT | 2 | GGTGCTCAATCACAGTTGGCAT |
| DENV3_13_LEFT | 1 | AGTGGAAGAAAGCAGAACTATAAGAGT |
| DENV3_13_RIGHT | 1 | TGGAGTTCACGTTCTCTGTCCA |
| DENV3_14_LEFT | 2 | GGAGAACCCTGGGAAGGAACAA |
| DENV3_14_RIGHT | 2 | ATGGCTGCCATTGAGGTATGTC |
| DENV3_15_LEFT | 1 | TGGACATCATATCTAGGAAAGACCAAAG |
| DENV3_15_RIGHT | 1 | AGGTTGCTCTGGAAGTGAGACC |

#### Supplementary Figure 1

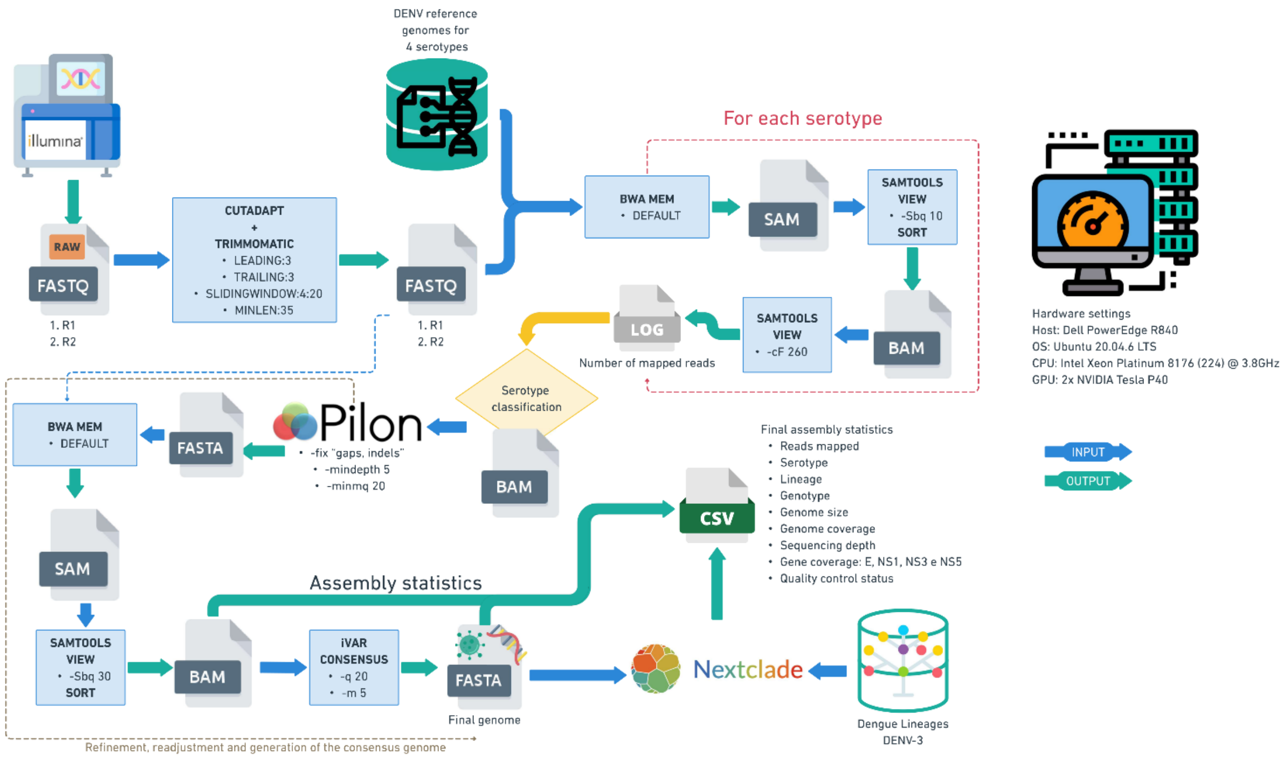

**Supplementary Figure 1. Workflow for the assembly, refinement, and classification of dengue virus sequences**. Paired end reads (R1, R2) undergo preprocessing to remove contaminants using Cutadapt and Trimmomatic. Filtered sequences are then mapped against reference genomes specific to each serotype using BWA-MEM. The resulting SAM files are converted to BAM and filtered with SAMtools to remove duplicates and unmapped reads. Assembly refinement is performed with Pilon, correcting indels and gaps, producing a more accurate final genome in FASTA format. After further filtering and consensus adjustments with iVAR, the consensus genome is generated. Data such as genome coverage and assembly statistics are compiled into a CSV file, including information on mapped reads, serotype, lineage, genome size, gene coverage, and overall assembly quality. Finally, the genome is classified using Nextclade for lineage determination.

#### Genome Assembly Pipeline

Initially, Cutadapt v2.8 (Martin, 2011) was used to remove adapters and primer sequences, following Trimmomatic v0.39 (Bolger et al., 2014) to filter out low-quality reads. Serotype identification was performed by mapping filtered reads to the reference genomes of the four dengue virus serotypes (DENV-1 to DENV-4: NC_001477.1, NC_001474.2, NC_001475.2, and NC_002640.1, respectively) using BWA-MEM v0.7.17-r1188 (Li, 2013), followed by counting reads in .bam file with SAMtools v1.12 (Li et al., 2009). The serotype was assigned based on the highest read count. The .bam file corresponding to the identified serotype was then refined with Pilon v1.24 (Walker et al., 2014), correcting gaps and indels using specified parameters, followed by a new mapping with filtered reads and assembly statistics collection. Finally, the consensus genome sequence in fasta format was generated using iVar v1.3.1 (Grubaugh et al., 2019) with parameters -q 20 and -m 5.

**Supplementary Table 2. Metadata of the 84 sequences used in the phylogeographic analysis, including unique sample ID, collection date, country, and state of origin.**

| ID | Collection Date | Country | State |
|---|---|---|---|
| EPI_ISL_19446000 | 2024-06-11 | Brazil | Pará |
| EPI_ISL_19446001 | 2024-05-14 | Brazil | Pará |
| CEVD_2401534 | 2024-08-21 | Brazil | Minas Gerais |
| CEVD_2401597 | 2024-10-30 | Brazil | Minas Gerais |
| CEVD_2401598 | 2024-10-25 | Brazil | Minas Gerais |
| CEVD_2401599 | 2024-09-30 | Brazil | Minas Gerais |
| CEVD_2401600 | 2024-09-20 | Brazil | Minas Gerais |
| CEVD_2401226 | 2024-01-24 | Brazil | Minas Gerais |
| CEVD_2401225 | 2024-03-04 | Brazil | Minas Gerais |
| CEVD_2401224 | 2024-03-06 | Brazil | Minas Gerais |
| CEVD_2401223 | 2024-03-26 | Brazil | Minas Gerais |
| EPI_ISL_17600856 | 2023-01-03 | Brazil | Roraima |
| EPI_ISL_17600857 | 2023-03-04 | Brazil | Roraima |
| EPI_ISL_17600858 | 2023-01-22 | Brazil | Roraima |
| EPI_ISL_17609844 | 2023-03-12 | Brazil | Suriname |
| EPI_ISL_18953953 | 2024-01-18 | Brazil | Minas Gerais |
| EPI_ISL_18953954 | 2024-01-19 | Brazil | Minas Gerais |
| EPI_ISL_18953955 | 2024-01-23 | Brazil | Minas Gerais |
| EPI_ISL_18953956 | 2024-01-24 | Brazil | Minas Gerais |
| EPI_ISL_18953957 | 2024-01-24 | Brazil | Minas Gerais |
| EPI_ISL_18953959 | 2024-01-25 | Brazil | Minas Gerais |
| EPI_ISL_18953960 | 2024-01-29 | Brazil | Minas Gerais |
| EPI_ISL_18542103 | 2023-11-09 | Brazil | São Paulo |
| EPI_ISL_18755964 | 2023-12-25 | Brazil | São Paulo |

| | | | |
|---|---|---|---|
| EPI_ISL_18871003 | 2023-11-28 | Brazil | São Paulo |
| EPI_ISL_18871004 | 2023-11-28 | Brazil | São Paulo |
| EPI_ISL_18871005 | 2023-12-04 | Brazil | São Paulo |
| EPI_ISL_18871006 | 2023-12-29 | Brazil | São Paulo |
| EPI_ISL_18871007 | 2023-12-28 | Brazil | São Paulo |
| EPI_ISL_19075352 | 2024-03-15 | Brazil | São Paulo |
| EPI_ISL_19179568 | 2024-03-13 | Brazil | São Paulo |
| EPI_ISL_19520624 | 2024-06-13 | Brazil | São Paulo |
| EPI_ISL_19520626 | 2024-04-01 | Brazil | São Paulo |
| EPI_ISL_19609374 | 2024-03-25 | Brazil | São Paulo |
| EPI_ISL_19609375 | 2024-04-03 | Brazil | São Paulo |
| EPI_ISL_19609376 | 2024-05-12 | Brazil | São Paulo |
| EPI_ISL_19609377 | 2024-10-16 | Brazil | São Paulo |
| OQ821509 | 2022-08-05 | Cuba | # |
| OQ821510 | 2022-08-06 | Cuba | # |
| OQ821521 | 2022-08-18 | Cuba | # |
| OQ821530 | 2022-08-24 | Cuba | # |
| OQ821531 | 2022-08-28 | Cuba | # |
| OQ821542 | 2022-09-13 | Cuba | # |
| OQ821552 | 2022-09-23 | Cuba | # |
| OQ821553 | 2022-09-23 | Cuba | # |
| OQ821554 | 2022-09-25 | Cuba | # |
| OQ821599 | 2022-12-22 | Cuba | # |
| OQ821613 | 2022-08-08 | USA | Florida |
| OQ821623 | 2022-12-23 | USA | Florida |
| OR150744 | 2022-12-19 | USA | Florida |
| OR150747 | 2022-12-23 | USA | Florida |
| OR150751 | 2023-01-09 | Cuba | # |
| OR150755 | 2023-01-08 | Cuba | # |
| OR574470 | 2023-09-01 | Italy | # |

| | | | |
|---|---|---|---|
| OR575180 | 2023-09-01 | Italy | # |
| OR654274 | 2023-05-28 | Cuba | # |
| OR771111 | 2023-01-25 | Cuba | # |
| OR771114 | 2023-02-21 | Cuba | # |
| OR771121 | 2023-03-29 | Cuba | # |
| OR771129 | 2023-07-13 | Cuba | # |
| OR771130 | 2023-07-03 | Cuba | # |
| OR771155 | 2023-08-20 | Cuba | # |
| OR771157 | 2023-08-22 | Cuba | # |
| OR771163 | 2023-08-29 | Cuba | # |
| OR771172 | 2023-09-06 | Mexico | # |
| OR771184 | 2023-09-11 | Dominican Republic | # |
| OR977070 | 2023-07-22 | Costa Rica | # |
| OR977095 | 2023-11-11 | Cuba | # |
| PP692453 | 2023-11-16 | Dominican Republic | # |
| PP692459 | 2023-12-02 | Puerto Rico | # |
| PP692464 | 2023-12-07 | Cuba | # |
| PP692471 | 2023-12-30 | Cuba | # |
| PP692472 | 2023-12-31 | Cuba | # |
| PP709303 | 2023-08-28 | Costa Rica | # |
| PP709304 | 2023-08-30 | Haiti | # |
| PP709308 | 2023-11-16 | Costa Rica | # |
| PP709326 | 2023-12-02 | Dominican Republic | # |
| PQ129528 | 2024-06-01 | USA | California |
| PQ129529 | 2024-06-04 | USA | California |
| PQ155005 | 2024-01-01 | Cuba | # |
| PQ155012 | 2024-03-25 | Cuba | # |
| PQ155038 | 2024-04-06 | Cuba | # |
| PQ155040 | 2024-04-13 | Puerto Rico | # |
| PQ155054 | 2024-06-06 | Puerto Rico | # |

*Note: Missing data indicated by #*

#### Results

#### Supplementary Figure 2

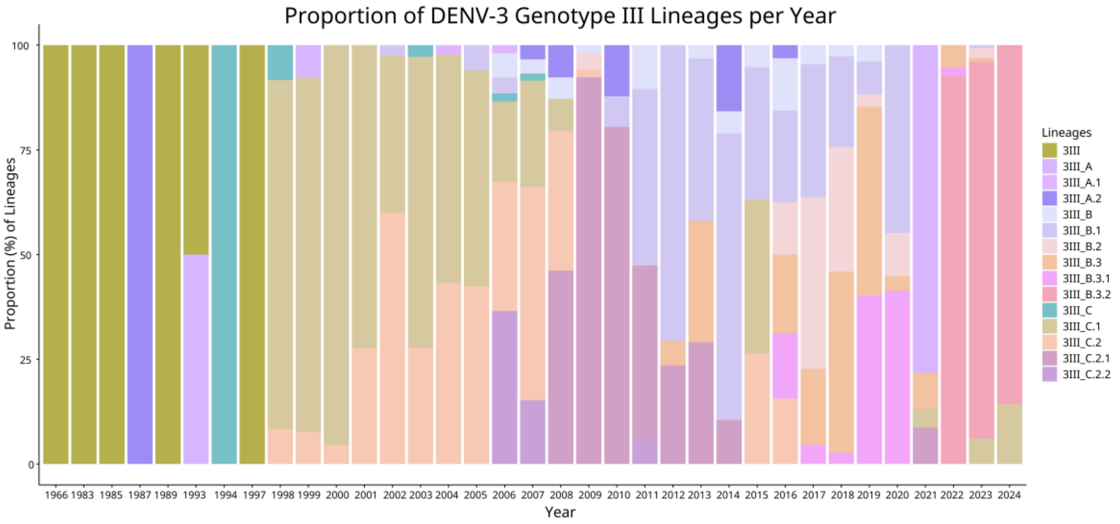

**Supplementary Figure 2. Proportional representation of DENV-3 genotype III lineages from 1966 to 2024.** Distinct lineages are represented by different colors, as indicated in the legend to the right. The Y-axis shows the percentage of each lineage relative to the total number of sequenced samples for the corresponding year (X-axis). These proportions reflect the availability of genomic data for each specific year.

#### Supplementary Figure 3

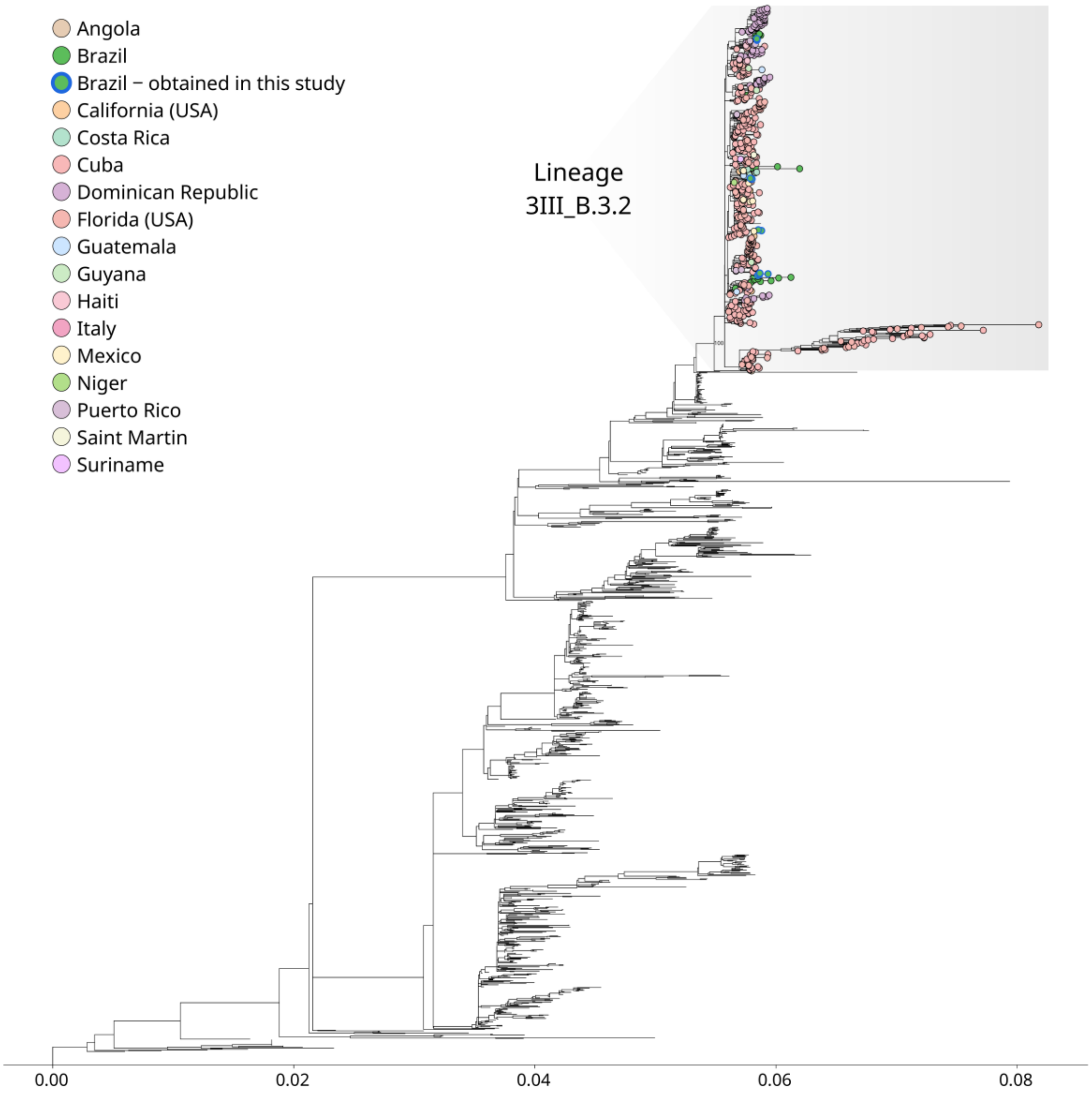

**Supplementary Figure 3. Maximum likelihood phylogenetic tree of DENV-3 genotype III.** The 3III_B.3.2 lineage is highlighted at the top of the tree (gray-shaded area). Each country is represented by a distinct color and the two new Brazilian genomes generated in this study are indicated by the blue outline, as indicated in the legend. Branch lengths represent genetic divergence, as shown by the scale. Ufboot support values for key branches in the 3III_B.3.2 lineage are also indicated on the tree.
